# Multiplexed Nanometric 3D Tracking of Microbeads using a FFT-Phasor Algorithm

**DOI:** 10.1101/763706

**Authors:** T. B. Brouwer, N. Hermans, J. van Noort

## Abstract

Many single-molecule biophysical techniques rely on nanometric tracking of microbeads to obtain quantitative information about the mechanical properties of biomolecules such as chromatin fibers. Their three-dimensional position can be resolved by holographic analysis of the diffraction pattern in wide-field imaging. Fitting this diffraction pattern to Lorentz Mie scattering theory yields the bead position with nanometer accuracy in three dimensions but is computationally expensive. Real-time multiplexed bead tracking therefore requires a more efficient tracking method. Here, we introduce 3D phasor tracking, a fast and robust bead tracking algorithm with nanometric localization accuracy in a *z-*range of over 10 µm. The algorithm is based on a 2D cross-correlation using Fast Fourier Transforms with computer-generated reference images, yielding a processing rate of up to 10.000 regions of interest per second. We implemented the technique in magnetic tweezers and tracked the 3D position of over 100 beads in real-time on a generic CPU. Its easy implementation, efficiency, and robustness can improve multiplexed biophysical bead tracking applications, especially where high throughput is required.

**Significance:** Microbeads are often used in biophysical single-molecule manipulation experiments and accurately tracking their position in 3 dimensions is key for quantitative analysis. Holographic imaging of these beads allows for multiplexing bead tracking but image analysis can be a limiting factor. Here we present a 3D tracking algorithm based on Fast Fourier Transforms that is fast, has nanometric precision, is robust against common artifacts and is accurate over 10’s of micrometers. We show its real-time application for magnetic tweezers based force spectroscopy on more than 100 chromatin fibers in parallel, but we anticipate that many other bead-based biophysical essays can benefit from this simple and robust 3 phasor algorithm.

## Introduction

Single-molecule techniques overcome ensemble averaging and can resolve unique and rare events at the molecular level [1]. By manipulation of microbeads, single-molecule force spectroscopy techniques revealed the mechanical properties of biomolecules such as DNA or RNA with unprecedented detail [2–5]. In addition, the interactions with proteins, like the DNA compaction by histones in eukaryotic chromatin [6–11] and prokaryotic architectural proteins [12–16], supercoiling [17–20], and repair processes [21–23] were extensively studied with magnetic tweezers (MT) or optical tweezers (OT), Acoustic Force Spectroscopy (AFS) [24–26] or tethered particle motion (TPM) [13, 27, 28]. These bead manipulation techniques have also been used to quantify the mechanical properties of other biological structures, such as extracellular protein collagen [29–31], or even entire cells [32].

The beads not only constitute a micron-sized handle to manipulate the molecules of interest, they also function as a label, whose position reflects the extension or deformation of the studied biomolecule. In OT, the position of one or two beads is generally measured from the deflection of a focused laser beam that is projected on a quadrant split detector [33, 34], yielding nanometric accuracy and kHz bandwidth in three dimensions. MT, TPM, and AFS however, generally use wide-field imaging with CCD or CMOS cameras and real-time image processing for position measurements. Next to sub-pixel accuracy, many applications require high framerates to resolve fast conformational changes or to capture the full spectrum of thermal motion for accurate force calibration [35]. Cameras with kHz frame rates or tens of megapixel resolution are currently available for fast or large field-of-view imaging [36, 37]. With such high-end hardware, real-time processing to resolve the three-dimensional position of the beads becomes rate-limiting. Moreover, multiplexing the image processing puts large demands on the processing power of the CPU. In some applications the GPU is employed to achieve sufficient speed [38–40].

Holographic imaging and subsequent fitting of the images to Lorentz Mie scattering theory (LMST) has been successfully used to convert movies of colloidal spheres into tracks of 3D coordinates [41]. Besides, one can accurately retrieve other physical characteristics that define the hologram, such as the bead radius and refractive index. Despite these advantages, bead tracking applications that are used in the single-molecule biophysics field generally use simpler, empirical methods to increase processing speed. A popular and fast method for bead tracking splits the tracking into 3 stages. First, the center of the bead is determined, either by computing the center of mass [42–44], or 1D or 2D cross-correlation with the mirrored intensity profile or predefined kernel [17, 34, 45–48]. Second, a radial intensity profile is computed. Third, this radial profile is compared to a previously calibrated look-up table (LUT) of radial profiles, and the *z*-coordinate is interpolated from the difference curve. Quadrant interpolation minimized crosstalk between *x, y*, and *z* coordinates, which significantly increased the tracking accuracy at the cost of being rather computationally intensive [49]. Cnossen *et al*. improved performance by shifting analysis to the GPU, which increased the speed and made it suitable multiplexed applications. This approach required specialized GPU hardware and advance software for analysis [50, 51].

Previously, we implemented the LUT bead tracking algorithm in our MT and used it to study the various transitions of chromatin fiber unfolding [6, 7, 52], as schematically depicted in Fig 1a. In these applications, we noticed that the dynamic range of the LUT method was sometimes insufficient, yielding an accuracy that depended on the bead height. As the composition of chromatin fibers may vary due to the quality of reconstitution [11], disassembly [6] or reflecting naturally occurring variations [53], we found it imperative to improve both the computation speed, allowing for tracking a large number of independent tethers, and the accuracy over a dynamic range that spans up to 10 micrometer. In our hands, the empirical LUT algorithm frequently flawed due to non-perfect imaging conditions, which led to discarding a large fraction of the beads, limiting the throughput. Tracking errors were enhanced by the increased field-of-view that is required for imaging multiple beads, which in practice yields more image artifacts, such as light gradients due to non-uniform illumination, astigmatism near the edges, or light obstructions by loose beads. Therefore, robustness against common image aberrations becomes increasingly important as the imaging settings can sometimes not be optimized for all beads.

**Figure 1.**
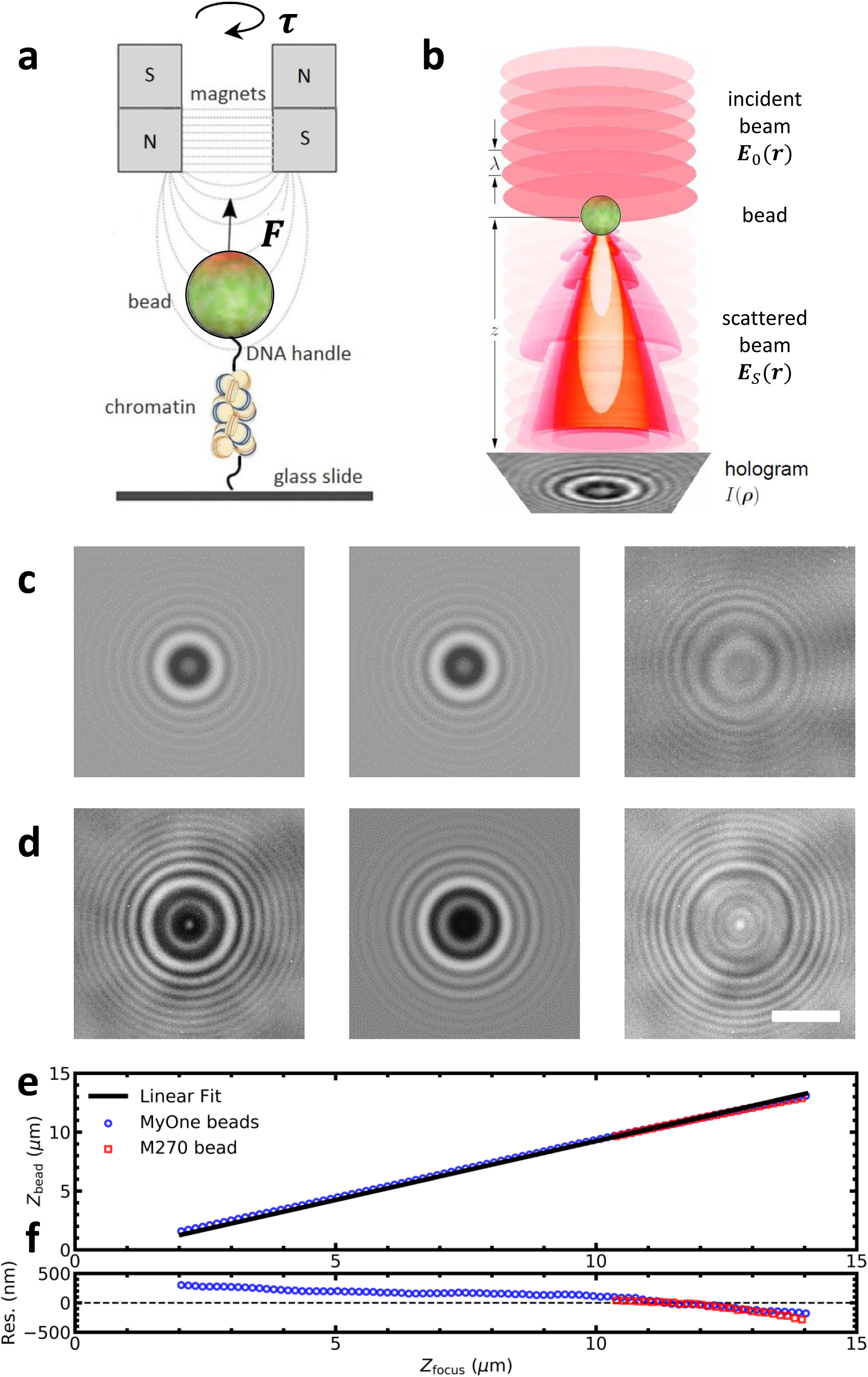
LMST cannot fully describe the holographic image of paramagnetic beads used in MT. a) Schematic drawing of a typical MT experiment. A molecule tethers a paramagnetic bead to the bottom of a flow cell. The bead is manipulated by a pair of magnets exerting force (***F***) and torque (***τ***). b) A holographic image *I*(***ρ***) is recorded originating from the interference of an incident beam ***E***_0_(***r***) with scattered the light ***E***_*s*_(***r***). The diffraction pattern is analyzed to obtain the three-dimensional position of the bead with nanometer accuracy. This image was adapted with permission from Lee *et al*. [41] © The Optical Society. c) The diffraction pattern (left) of a 1.0 µm diameter paramagnetic bead (Dynabeads MyOne Streptavidin T1, *Thermo Fisher Scientific*) was fitted with LMST (center). The fit to equation 10 yielded x = −70±1 nm, y = 33±1 nm, z = 8300±100 nm, *n*_bead_ = 1.9±0.1, *α* = 0.9±0.1, *β* = 57±1, *γ* = 57±1 (fit±standard error). The values *a* = 0.5 μm and *n*_medium_= 1.33 were fixed. The residual image (right) shows that some features could not be reproduced by LMST. d) The diffraction pattern of a 2.8 µm diameter paramagnetic bead (Dynabeads M270 Streptavidin, *Thermo Fisher Scientific*) yielded a worse fit: x = −162 ± 1 nm, y = 84 ± 1 nm, z = 9500±100 nm, n_bead_ = 1.8±0.1, *α* = 0.7 ± 0.1, *β* = 49 ±1, *γ* = 67 ± 1. The values *a* = 1.4 μm and *n*_medium_= 1.33 were fixed. Scale bar: 3 µm. e) Two paramagnetic beads (MyOne in blue and M270 in red) were moved through focus, and the recorded holographic movies were fitted to LMST. For clarity, every fifth data point was plotted. The fits of the diffraction pattern (black lines) did not converge close to focus. Sufficiently far from the focus, the obtained bead height was proportional to *z*_focus_. f) Residual of the linear fit of the bead height as a function of the focus height.

Using the power of 2D Fast Fourier Transforms (FFT) to compute cross-correlations with computer-generated reference images, we reduced the computational effort to three FFTs and skipped the generation and comparisons with radial profiles, which is the computationally most expensive part of traditional bead tracking. Instead, translations in the *z*-direction were captured into a single parameter, the phase, that we show to be proportional to the height. This new tracking algorithm, which we call 3D phasor tracking (3DPT), is simple, fast and robust and meets all criteria for real-time multiplexed nanometric bead tracking experiments on a generic CPU.

## Materials & Methods

### Magnetic tweezers setup

A multiplexed MT setup equipped with a NIKON CFI Plan Fluor objective MRH01401 (NA = 1.3, 40x, Oil, *NIKON Corporation*, Tokyo, Japan) was used to track paramagnetic beads. Samples were measured in custom-built flow cells mounted on a multi-axis piezo scanner P-517.3CL (*Physik Instrumente GmbH & Co. KG*, Karlsruhe, Germany). A field-of-view of 0.5×0.5 mm^2^ was captured on 25 Mpix Condor camera (cmv5012-F30-S-M-P8, *CMOS Vision GmbH*, Schaffhausen, Switzerland) using an infinity-corrected tube lens ITL200 (*Thorlabs*, Newton, USA). The camera was read out by a PCIe-1433 frame grabber (*National Instruments*, Austin, USA) integrated with a T7610 PC (*Dell*, Round Rock, USA) equipped with a 10-core Intel Xeon 2.8 GHz processor (E5-2680 v2, *Intel*, Santa Clara, USA) and 32GB DDR3 memory. The setup measured the full frame at 30 frames per second. Each pixel measured 112 nm in this configuration. The flow cell was illuminated with a 100 mW 645 nm LED-collimator-packaged (LED-1115-ELC-645-29-2, *IMM Photonics GmbH*, Unterschleißheim, Germany).

### Software

All MT control software, LMST fitting tools, and tracking software were written in LabVIEW 2014 (*National Instruments*, Austin, USA) and are available upon request.

### Spectral analysis of tracking accuracy

A solution 1 pg/µl 2.8 µm diameter paramagnetic beads was deposited onto a cover slide and heated to 95 °C for several minutes to melt the beads to the glass. Subsequently, the cover slide with immobilized beads was mounted into the flow cell and placed onto the setup. Immobilized beads were tracked for 120 seconds. To obtain *σ*^2^/*f*_*s*_, the PSD was calculated and fitted with a horizontal line for *f* > 5 Hz.

### Lorentz Mie scattering theory

LMST fitting was implemented following [41, 54, 55]. The diffraction pattern *I*_*LMST*_(*ρ*) results from the interference between the incident field *E*_0_(*r*) and the field scattered off the particle *E*_*s*_(*r*):

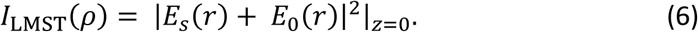

The incident field was described by:

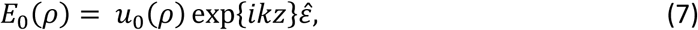

where the incident field was uniformly polarized in the 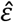-direction so that amplitude *u*_0_(*ρ*) at position *ρ* = (*x, y*) in plane *z* = *z*_*p*_ of the particle is equal to that in the focal plane *z* = 0. The wavenumber of the propagating wave was *k* = 2*πn*_*m*_/*λ*, where *n*_*m*_ was the refractive index of the medium and *λ* was the wavelength of the light in vacuum.

The scattered field was described by:

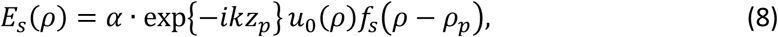

where *f*_*s*_(*ρ*) was the LMST function which depended on bead radius *a, n*_*p*_, *n*_*m*_, and *λ. α* ≈ 1 and accounted for variations in the illumination.

The diffraction pattern was normalized in the *z* = 0 plane by *β*:

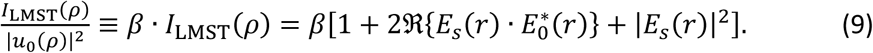

The normalized image scaled with the calculated Mie scattering pattern *f*_*s*_(*r*) by:

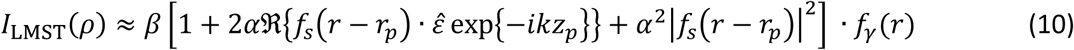

where *f*_*γ*_(*r*) is a Hamming filter of width *γ* which represented the decay of the diffraction pattern from the center due to the limited spatial coherence of the light source:

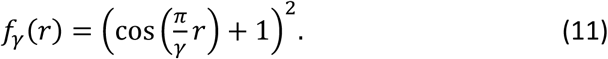

Fitting equation 10 yielded the physical parameters *a, n*_*p*_, *n*_*m*_, *x, y, z*, and scaling coefficients *α, β*, and *γ*. The three-dimensional position and radius *a* of the bead was typically fitted with nanometer precision. The refractive index *n*_*p*_ was reproducible between beads within one part in a thousand [41].

### 3DPT robustness simulations

We simulated movies of beads randomly moving in three dimensions using LMST using the following parameters: *a* = 0.5 μm, *n*_*p*_ = 1.9, *n*_*m*_ = 1.33, *α* = 1.0, *β* = 54, *γ* = 45, and *λ* = 645 nm. This approach yielded realistic diffraction patterns of 1.0 μm diameter paramagnetic beads. Parameters *n*_*p*_, *α, β*, and *γ* were average values obtained from fitting the experimental patterns of 186 separate beads (data not shown). For these simulations, the phase was calibrated using 15 polynomials and 15 reference periods. For every simulation, we computed 3600 holograms (150×150 pixels) of beads randomly moving in three dimensions (*dx, dy* = 0 ± 5 nm, *dz* = 12000 ± 5 nm). The simulations were equivalent to a 120 seconds measurement on a 30 Hz camera. The simulated beads were tracked using 3DPT, and the time traces were converted to PSD to extract *σ*^2^/*f*_*s*_, similar to experimental data.

Five common aberrations were superimposed on the simulated diffraction patterns. Poisson noise was added to the diffraction pattern. Interlacing was simulated by multiplying every other row of pixels with a gain factor. 100% interlacing corresponded to a gain of 2. Light gradients were simulated by adding a slope in the *y*-direction of the diffraction pattern. 100% light gradients corresponded to a curve which rose to a maximum background intensity of 255. Astigmatism was simulated by resampling the columns in the *y*-direction over a smaller number of pixels. 100% astigmatism corresponded to an aspect ratio of 2. Shift out of the ROI was simulated by moving the bead in the *x*-direction until the average bead center was shifted by 50% of the ROI.

### Characterization of mechanical vibrations and drift

To characterize 3DPT accuracy experimentally, time traces of 10 immobilized beads were recorded simultaneously. The effects of mechanical vibrations and drift, were largely removed by averaging these traces and subtracting it after applying a 1 Hz low-pass filter. The standard deviations of the resulting time trace yield the experimental tracking accuracy.

### Quantification of the step size of the unwrapping of DNA from the histone core

The length of the stepwise unwrapping of DNA from native nucleosome cores was measured using a method developed by Kaczmarczyk, described in detail in [56]. In short: each data point in the force-extension curve was compared to the theoretical extension of a given a contour length of free DNA, following a Worm Like Chain. The theoretical standard deviation for each point was computed using equipartition theorem and the derivative of the force-extension relation of the wormlike chain and supplemented by the tracking error. Next, the z-score and corresponding probability that the data point belonged to this contour length was calculated. The probabilities for all data points at given contour length were summed and this procedure was iterated for all contour lengths between 0 and the contour length of the DNA substrate. Peaks in the plot of the summed probability as a function of contour length were attributed to a stable state of unfolding of the chromatin fibers and distances between neighboring peaks reflect single unwrapping events.

## Results

### Tracking of super-paramagnetic beads using Lorenz-Mie scattering theory

The scattering of light by colloidal particles and its interference with the incident light is described in LMST. Fig 1b depicts the contrast mechanism of holographic imaging of a colloidal bead (adapted from [41]). The interference of the incident beam ***E***_**0**_(***r***) with the light that is scattered off the bead ***E***_***s***_(***r***), yields a circularly symmetric hologram *I*(***ρ***) in the image plane. The center of the hologram corresponds to the *xy*-position of the bead. As the image plane moves away from the location of the bead, the interference pattern expands, leading to more and larger rings around the center of the bead.

We used LMST to fit the 6 parameters that describe the diffraction pattern: *x, y, z*, bead radius *a*, refractive index of the medium *n*_*m*_, refractive index of the bead *n*_*p*_, and three additional scaling parameters *α, β, γ* [54, 55] (*see* Materials & Methods). We tested two types of super-paramagnetic beads, either with a diameter of 1 µm (Dynabeads MyOne Streptavidin T1, *Thermo Fisher Scientific*, Fig 1c, left) or 2.8 µm (Dynabeads M270 Streptavidin, *Thermo Fisher Scientific*, Fig 1d, left), which are commonly used in MT. In both cases, we obtained a reasonable fit (Fig 1c and d, center), though spherical shapes in the residual images (Fig 1c and d right) indicate that there is a systematic discrepancy between the LMST fit and the experimentally obtained holographic images. The residuals were generally larger for the 2.8 µm beads than for the 1 µm. Though we did not further investigate this difference, we attribute it to the mixed composition of these beads, which may not fully be captured in a single refraction index used in LMST.

For MT force spectroscopy applications, the *z*-coordinate is the most important parameter that can be extracted for the hologram, as it quantifies the extension of the tether. In Fig 1e, the fitted *z*-coordinate is plotted, when a bead that was fixed on a cover slide was moved linearly through the focus using a piezo stage. Fitting the obtained diffraction patterns with LMST at corresponding height from focus, a fairly accurate position could be obtained over a range of more than 10 µm for the smaller and about 5 µm for the larger bead. The *xy*-coordinates also yielded reproducible results (data not shown). A closer look at the residual of fitting a linear curve to the measured versus applied height however, Fig 1f, shows that the systematic errors in the *z*-direction typically amounted hundreds of nanometers for our paramagnetic beads. The relative error can be smaller for smaller ranges, which makes it still useful for small tethers.

The main limitation of LMST fitting however, is the processing speed. Fitting the diffraction pattern of a single bead in a 100×100 pixels region-of-interest (ROI) took several seconds on our CPU. For off-line applications this may not be problematic, although implementation of off-line multiplexed bead tracking would imply storage and processing of very large data files. Real-time processing has the advantage that the experimenter can rapidly asses the quality of the measurements or make adjustments during the measurement. LMST fitting cannot achieve real-time processing with current computing power, which requires a more efficient tracking method.

### 3D Phasor Tracking

Here we introduce a novel method called 3D phasor tracking (3DPT), which exploits the circular symmetry of the LMST diffraction pattern, and its gradual expansion when the bead moves in the *z*-direction, to compute bead coordinates. Each step of the tracking algorithm is depicted in Fig 2. We calculated the *xy*-coordinate by cross-correlating the ROI with a computer-generated reference image, rather than its mirrored image. The complex reference image *I*_*n*_(*r*) resembles the holographic image, but consists of a single spatial frequency, characterized by period *k*_*n*_:

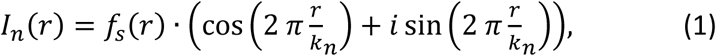

with *r* the distance from the center of the ROI of size *s*. The reference image is spatially filtered with filter *f*_*s*_(*r*):

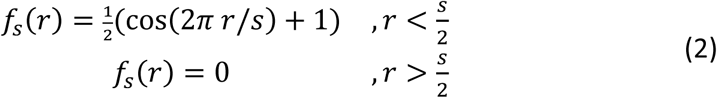

**Figure 2.**
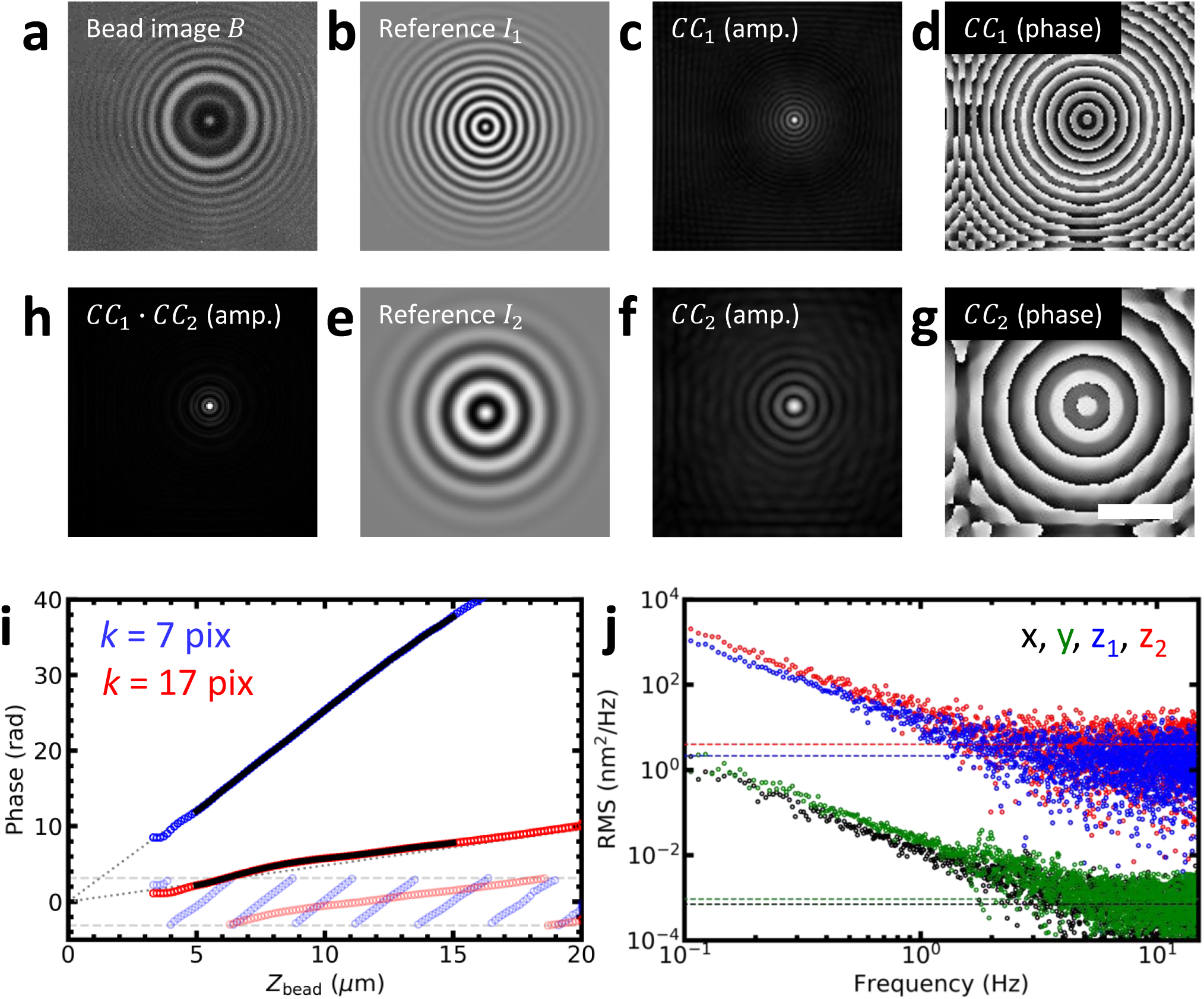
The principle of 3D phasor tracking (3DPT). The holographic image (a) was cross-correlated with two complex reference images *I*_1_ (b) and *I*_2_ (e), computed using equation 1 for period *k*_1_ = 7 pix and *k*_2_ = 16 pix. The cross-correlation yielded two complex images *CC*_1_ and *CC*_2_, displayed as amplitude (c, f) and phase (d, g). The amplitudes of *CC*_1_ and *CC*_2_ were multiplied, resulting in a sharp peak at the *xy*-position of the bead (h). The phases at the peak, *φ*_1_ and *φ*_2_, scaled approximately linearly with bead height *z*. Scale bar: 3 µm. i) The relation between *φ* and *z* was calibrated *a priori* using a measurement in which the focus is linearly shifted in time resulting in phases *φ*_1_ (semi-transparent blue circles) and *φ*_2_ (semi-transparent red circles). Subsequently, *φ*_1_ and *φ*_2_were phase-unwrapped, eliminating 2π phase jumps (blue and red circles beyond the horizontal dashed lines). A linear increase in phase, starting at focus roughly describes the phase-height relation (grey dotted lines). The unwrapped phases *φ*_1_(*z*) and *φ*_2_(*z*) were fitted with a polynomial, starting 5 µm above focus (black lines). For clarity, every fifth data point was plotted. j) The tracking accuracy of 3DPT was calculated by spectral analysis, yielding an accuracy of 4 ± 1 nm^2^/Hz for *z*_1_, and 2 ± 1 nm^2^/Hz for *z*_2_. The accuracy in *x* and *y* was several orders of magnitude smaller: 0.7 · 10^−3^ ± 0.1 nm^2^/Hz for *x*, and 0.9 · 10^−3^ ± 0.1 nm^2^/Hz for *y*.

Two examples of reference images for two different periods are shown in Fig 2b and 2e.

The experimentally measured diffraction pattern of the bead *I*_bead_ is cross-correlated with the reference images yielding cross-correlation *CC*_*n*_:

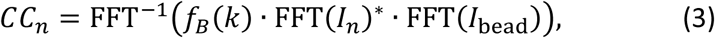

where a band-pass filter *f*_*B*_(*k*) is used in the frequency domain:

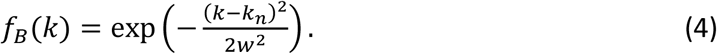

The computer-generated images of the reference signal and its filters are depicted in Fig S1.

Figs 2c-g show the amplitude and phase images for of the cross-correlation with a typical hologram, shown in Fig 2a, for periods *k*_1_ = 7 pixels and *k*_2_ = 16 pixels. The amplitude image of the *CC*_*n*_ in both cases featured a single peak that represents the shift of the bead relative to the center of the image. Two amplitude images obtained by cross-correlating with reference images with two different periods were multiplied yielding a sharper peak at the *xy*-position of the bead (Fig 2h), which was measured with sub-pixel accuracy using polynomial interpolation in 2D.

The *z*-position of the bead was obtained from the phase *φ*_*n*_ at the *xy*-position of the bead. The phase images featured concentric rings, around the bead center. In Fig 2i, we plotted the phase at the bead center of a fixed bead that was moved through the focus in the *z*-direction. Several micrometers above the focus, we obtained good cross-correlations with distinct peaks at the bead position. The phase increased proportional to the bead height but was wrapped between -π and π. Unwrapping of the phase was trivial since the height of the bead increased linearly in time. The phase signal could thus be unwrapped unambiguously as shown in Fig. 2i. We quantified this curve by fitting a polynomial function to the experimental *φ*_*n*_(*z*) data, which was inverted to compute the bead height as a function of the measured phase, *z*(*φ*_*n*_), during subsequent experiments.

Far from focus, *φ*_*n*_ increased, to a good approximation, linearly with *z*. Empirically, we found for sufficiently defocused beads:

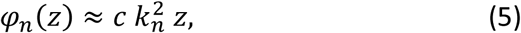

in which *c* represents a calibration factor that depends on the magnification and the refraction index of the immersion medium, and did not change between experiments or bead sizes. This linear approximation is plotted in Fig 2i as dotted lines, and it can be seen that deviations become larger near focus and for larger periods, as near-field effects of the light scattering become more prominent.

In tracking experiments in which there is no prior knowledge of the bead position, a single phase cannot be converted unambiguously into a unique *z*-position due to phase wrapping. A common solution to this phase unwrapping problem is the use of multiple frequencies [57–60], which was implemented by computing at least two *CC*_*n*_ images with different periods in the reference images. Provisional *z*-coordinates, which corresponded to the expected *z*-range of the bead, spaced by 2π in phase, were calculated. The set of *z*-coordinates corresponding to different spatial frequencies that showed the smallest variation was then selected and averaged to compute the final *z*-coordinate. Thus, by computing two, or more, reference images and subsequent cross-correlation with an experimental holographic image, it was possible to determine the three-dimensional position of the bead unambiguously.

### Performance of 3DTP

The performance of the novel 3DPT method was tested in multiple ways. First, the power spectral density (PSD) of a time trace of an immobilized bead was computed (*see* Materials & Methods), as shown in Fig 2j. The tracking accuracy, expressed in *σ*^2^/*f*_*s*_, was determined as the plateau value of the PSD at frequencies over 2 Hz [33], since thermal drift and mechanical vibrations introduced low-frequency fluctuations that resulted in increased amplitudes at smaller frequencies (1/*f* noise). The reference image with the highest spatial frequency (*k* = 7 pix) performed best: *σ*_*z*_^2^/*f* = 2 nm^2^/Hz. For the lower spatial frequency (*k* = 16 pix) we obtained an accuracy of *σ*_*z*_^2^/*f* = 4 nm^2^/Hz. The tracking accuracy in the *x*- and *y*-direction was several orders of magnitude higher: *σ*_*x*_^2^/*f* = 0.7 · 10^−3^ nm^2^/Hz and *σ*_*y*_^2^/*f* = 0.9 · 10^−3^ nm^2^/Hz. Thus, for a typical frame rate of 30 Hz, we can expect a tracking accuracy of 0.2, 0.2 and 10 nm for the *x, y* and *z*-coordinate after cross-correlation with a single reference image.

Next, to illustrate the increased performance with an increased number of reference images, we plotted the phase calibration graphs obtained for four reference images, shown in Fig 3. The *φ*_*n*_(*z*) curve converged to a single point following equation 5, which allowed us to unequivocally assign the focus offset. The deviations from linearity, starting 5 µm above focus, were fitted with a 5^th^ order polynomial over a range of 10 µm. The residuals of the fits give a good estimation of the dynamic range: the standard deviation, *σ*_res_, was generally below 15 nm, as depicted in Fig 3b. We systematically tested the dependence of the accuracy of the 3DTP method by evaluating the standard deviation of the residuals as a function of the reference frequency. The lowest *σ*_res_ was found for *k* = 10 pix. For *k* < 7 pix we did not obtain distinct correlation peaks, reflecting the diffraction-limited character of the holographic images. For *k* > 10 pix, *σ*_res_ gradually increased up to 15 nm, which is still rather accurate for a dynamic range of 10 µm (Fig 3c). As could be expected from visual inspection of the holographic images, the further the bead was defocused, the less high-frequency information was obtained. This was reflected by the amplitudes of the cross-correlations, that were plotted in Fig 3d. Whereas for *k* = 7 pix, the highest cross-correlation was obtained 10 µm above focus and vanished at 20 µm, larger periods peaked further from the focus. From Fig 3a-d it is clear that the dynamic range is not limited to 10 µm.

**Figure 3.**
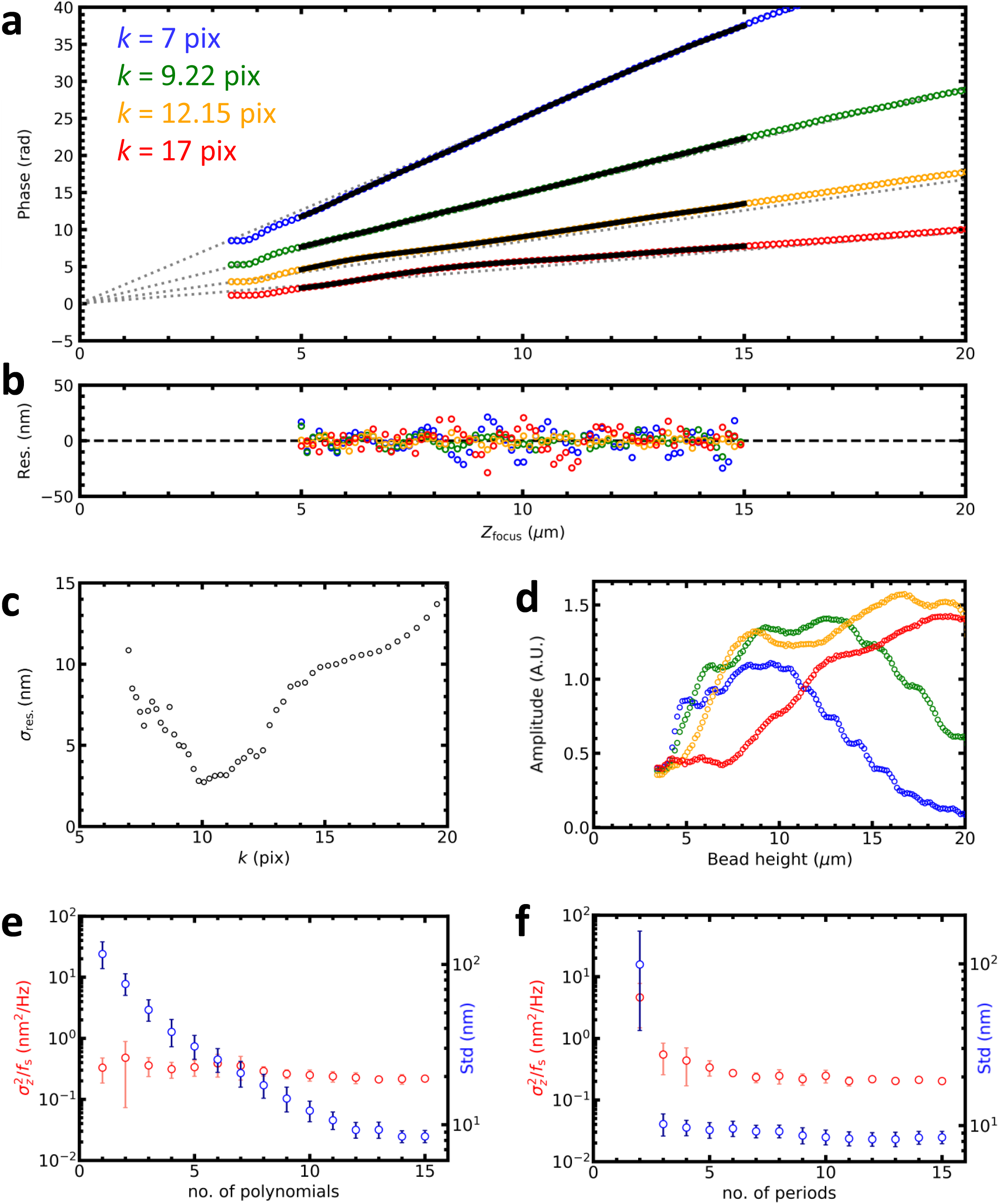
Multiple reference images increase the tracking accuracy. a) Phase calibration with 4 reference images (colored circles). *k*_*n*_ was logarithmically sampled between 7 and 16 pixels. The curves were fitted with a 5^th^ order polynomial over a range of 10 µm (black lines). All curves converged in focus and approximated a straight line (grey dotted lines). b) The residuals of the polynomial fit for each reference image (colored circles). c) The tracking accuracy obtained with a single reference image varied with its period. The best accuracy was obtained for *k*≈10 pix. Below *k*=7 pix, the correlation did not yield a distinct peak. d) The amplitude of the cross-correlation varied with *z* and *k*. Larger periods were more prominent as the bead was shifted further out of focus. For clarity, every fifth point was plotted in graph a, b, and d. e) The tracking accuracy in the *z*-direction (red dots) slightly increased with the number of polynomials used during phase calibration. The errors significantly reduced up to approximately 5 polynomials. The average standard deviation of the residuals of the polynomial fit (blue dots) linearly decreased up to approximately 10 polynomials. f) The tracking accuracy in the *z*-direction (red dots) increased significantly with the number of reference images, up to approximately 10 reference images. Especially the use of 3 instead of 2 reference images was effective: approximately a 10-fold increase in accuracy was established. 15 polynomials were used during phase calibration. The average standard deviation of the residuals of the polynomial fit (blue dots) was not affected by the number of reference images when 3 or more images were used. The errors depicted in Fig 3e and 3f indicated the standard deviation of 6 independent beads.

The periodic modulation in Fig 3b suggests that better accuracy could be obtained by fitting higher order polynomials. Increasing the order of the polynomials did decrease the average standard deviation of the residuals of the polynomial fit, although this was not reflected in the PSD analysis of the experimental data, shown in Fig 3e. This is probably due to the slow fluctuations that remain in the residual of the polynomial fit, which would be represented in the low-frequency part of the power spectrum.

The tracking accuracy in the *z*-direction could be increased by combining more reference images, at the cost of increased computational time. Fig 3f shows that the accuracy, as quantified from the PSD, increased about 10-fold when more reference images were used. The accuracy converged to *σ*_*z*_^2^/*f* = 0.2 nm^2^/Hz for *n* = 5, implying 2.4 nm accuracy for 30 Hz imaging or 1 nm at 5 Hz. The average standard deviation of the residuals of the polynomial fit decreased similarly. For accuracy in the *x*-and *y*-direction, it was sufficient to use more than 2 reference images to achieve sub-nm accuracy at 30 Hz.

The computation speed was up to 10.000 diffraction patterns per second for 100×100 pix ROIs, even when 4 reference images were used. Note that the FFT of the reference images can be done prior to the tracking, so only (*n* + 1) FFTs need to be computed in real-time for *n* reference images. Subsequent processing, i.e. calculating the *xy*-position, computing the *z*-coordinate from the calculated phase and unwrapping the phase, was much faster. Thus, 3DPT is a versatile, accurate and fast method for holographic tracking of microbeads with nanometer accuracy.

### The robustness of 3DTP

To evaluate the robustness of our tracking algorithm we simulated test data using LMST for 1.0 µm diameter beads that were moving randomly in three dimensions. We simulated the diffraction patterns 12 μm above focus, where the accuracy of LMST was the highest (Fig 1f). This approach allowed us to systematically introduce distortions and to compare tracking results with the known coordinates that were inserted into the LMST.

Fig 4a shows a simulated image in which we have introduced Poisson noise, representing the signal to noise of a typical camera. While the accuracy of the *x*-and *y*-coordinates was hardly affected and remained constant at 0.5 nm^2^/Hz, the accuracy in the *z*-direction decreased from 0.05 to 10 nm^2^/Hz when the noise increased for 0 to 20 greyscale units in an 8-bit image. Typical experimental noise intensities (∼3 greyscale units) resulted in 1 nm^2^/Hz accuracy in the *z*-direction, close to the experimentally obtained accuracy in figure 3e-f.

**Figure 4.**
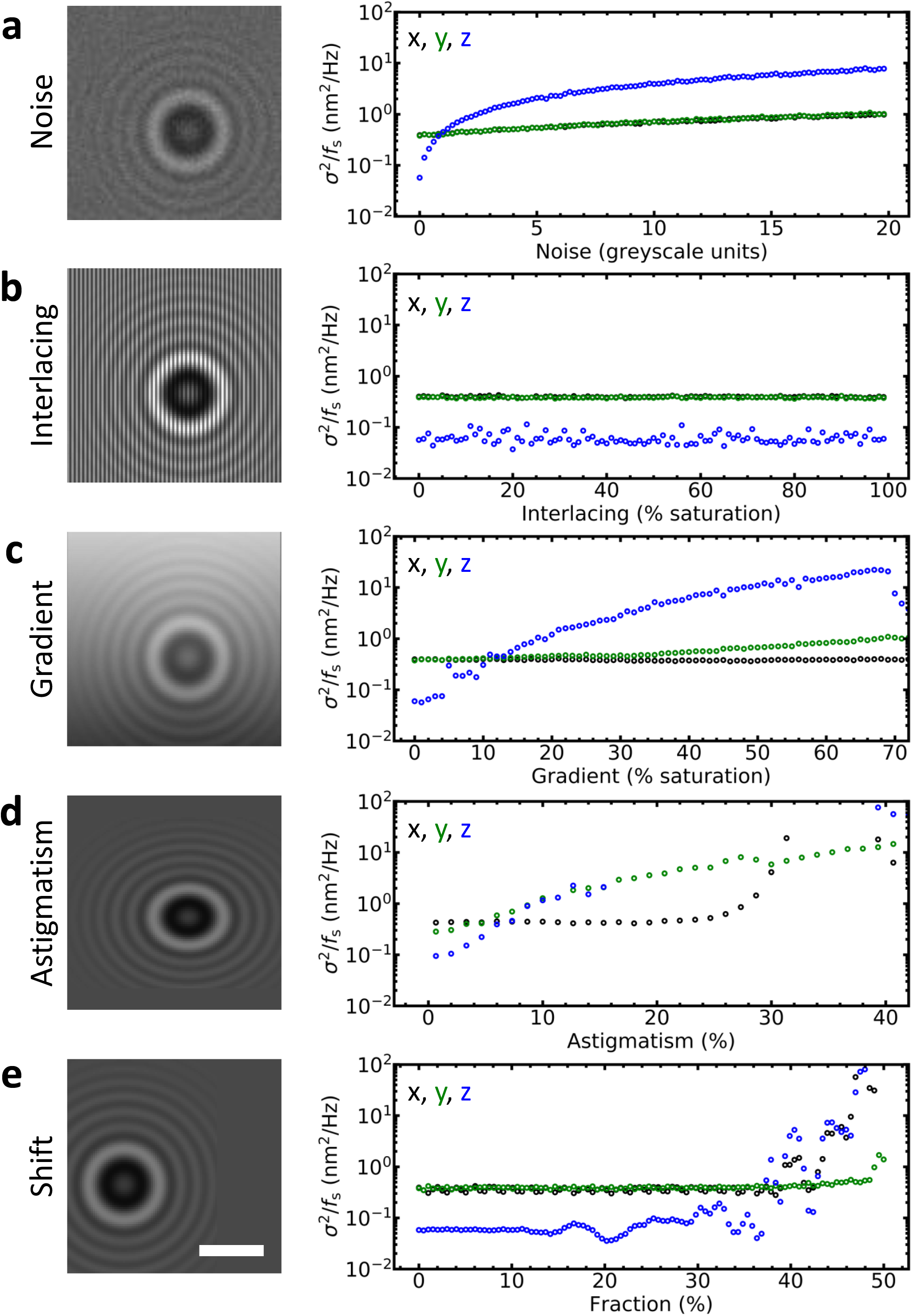
3DPT is robust against image aberrations. To evaluate the robustness of 3DPT, the diffraction patterns of 1.0 µm diameter paramagnetic beads were simulated with LMST. In the absence of image artifacts, the tracking accuracy of simulated images was 60 · 10^−3^ nm^2^/Hz in the *z*-direction. The diffraction patterns were superimposed with aberrations typically observed in MT and other bead tracking techniques. Note that the amplitude of the PSD remained below 100 nm^2^/Hz, corresponding to 55 nm (0.5 pix) at a frame rate of 30 Hz, in all cases except for astigmatism exceeding 30%. Typical experimental values for our microscope were ∼3 greyscale units for Poisson noise, 0% for interlacing, ∼15% for light gradients, ∼2% for astigmatism, and ∼5% for shift. a) Poisson image noise mainly affected the accuracy of the *z*-coordinate and resulted in a 10-fold decrease in accuracy for an amplitude of 20 greyscale units. The tracking accuracy in the *x*-and *y*-direction only decreased a factor of 2. b) Interlacing did not affect the tracking accuracy. c) A light gradient in the ROI up to 20% still yielded an accuracy in the *z*-direction below 1 nm^2^/Hz. The tracking accuracy was only slightly affected in the *y*-direction, the direction of the light gradient, while the *x*-direction was unaffected. When the light gradient exceeded 70% no correlation peak was found. d) Astigmatism significantly affected tracking accuracy in three dimensions. e) Our tracking method was unaffected when the bead center was shifted less than 30% out of the ROI. Exceeding a shift of approximately 40% resulted in the complete loss of tracking in both the *z*-direction and the *x*-direction (the direction that the bead was moving). Scale bar: 3 µm.

The robustness of the algorithm was further tested in Figs 4b-e, in which we simulated several other distortions that are frequently observed in experimental imaging. Interlacing, which was prominent in analog CCD cameras, did not affect the tracking accuracy (Fig 4b). Distortions with lower spatial frequencies, such as light gradients, did decrease the accuracy of the calculated *z*-coordinate (Fig 4c). Astigmatism up to 15% resulted in moderate increases of the error (Fig 4d). Larger astigmatism was more problematic. However, such distortions should not occur in properly designed microscopes.

We also quantified the robustness of the tracking algorithm when the diffraction pattern was not fully contained within the ROI. Since there is always a trade-off between the size of the analyzed ROI and computation speed, it is in the interest of increasing throughput to reduce the ROI as much as possible. Moreover, restricting the ROI reduces the probability that other beads enter the ROI, which hampers accurate tracking. We observed that tracking accuracy was not affected until the bead was shifted by more than 30% out of the ROI (Fig 4e). In conclusion, our simulations showed that 3DPT was robust against many types of typical aberrations.

Finally, the performance of 3DPT was demonstrated experimentally by simultaneous measurement of the three-dimensional position of 10 immobilized 2.8 µm diameter paramagnetic beads. Drift and mechanical vibrations, which dominated our measurements, as reflected in the power spectral densities (Fig 2j), were characterized, isolated, and removed following the approach described in Materials & Methods. A representative time trace in three dimensions was shown in Fig 5a-c, yielded a standard deviation in the *x*-and the *y*-direction of 0.3 nm and in the *z*-direction 1.6 nm, roughly matching the simulated values.

**Figure 5.**
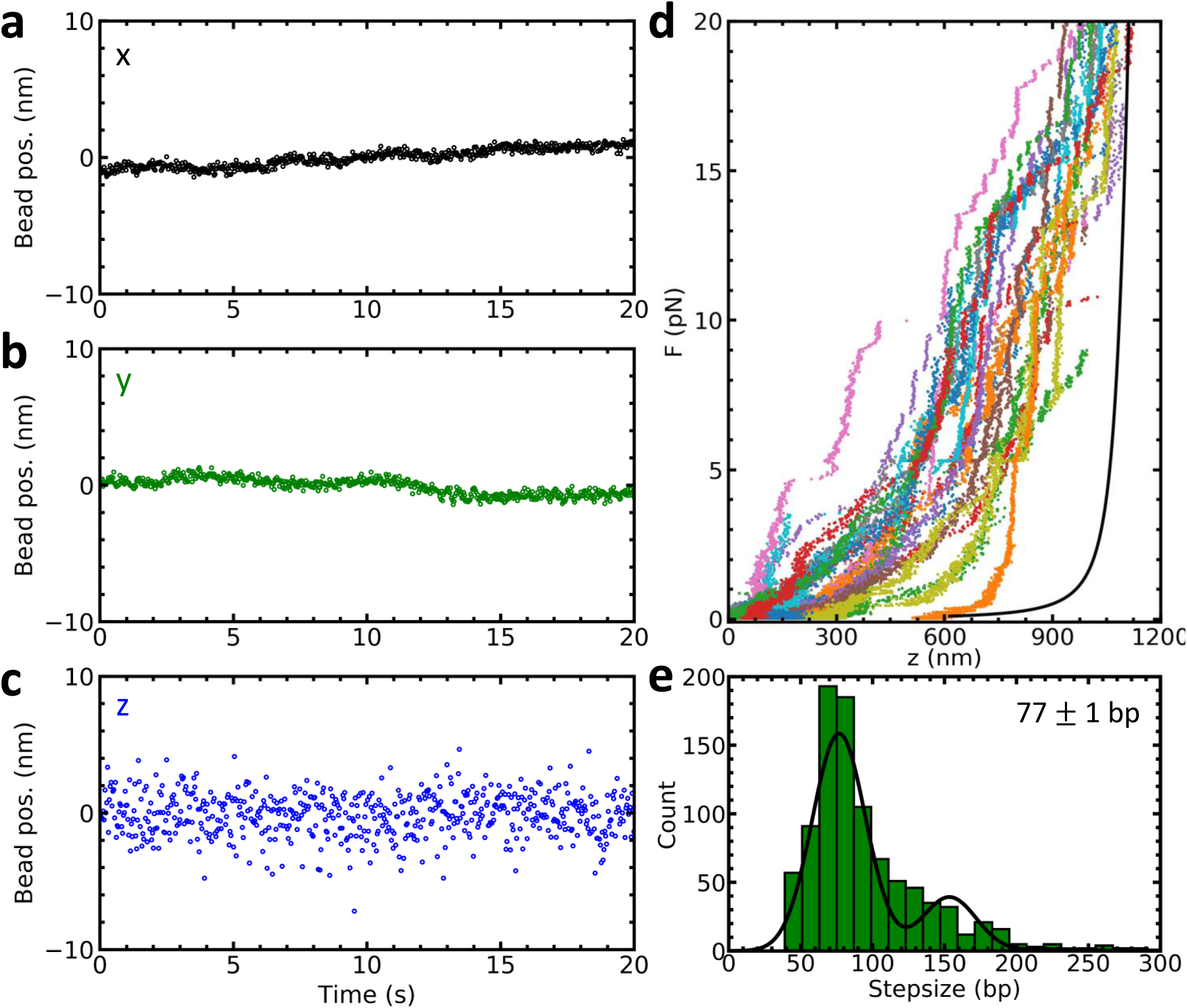
Simultaneous nanometric tracking of multiple paramagnetic beads yields nanometer accuracy in three dimensions. a-c) 10 immobilized 2.8 µm diameter paramagnetic beads (Dynabeads M270 Streptavidin, *Thermo Fisher Scientific*) were recorded for 20 seconds and analyzed using 3DPT using 15 polynomials during phase calibration and 15 reference images during tracking. Drift and mechanical vibrations were characterized, isolated, and removed following the approach described in Materials & Methods, and a typical trace was plotted in the x-(a), y-(b), and z-direction (c). Some residual drift that was present in all dimensions indicated imperfect immobilization of the beads. The distribution of the coordinates yielded a standard deviation of *σ*_*xy*_ = 0.6 nm in the *x*-and the *y*-direction and *σ*_*Z*_ = 1.6 nm in the *z*-direction. d) 25 native chromatin molecules, reconstituted *in vivo* [53], were stretched and unfolded with magnetic tweezers. Although the complexes were heterogeneous, they unfolded in three distinct transitions [63]. The last transition, where the last singe turn of DNA unwraps from the histone core, takes place above +/-5 pN (for native chromatin) and is recognized by its stepwise nature. e) The step size distribution was plotted in a histogram (bin width = 12 bp) and fitted with a triple Gaussian. The histogram contained data from 111 curves from which a selection was plotted in figure 5b. The step size was easily resolved with our approach, and measured 77±1 bp, taking into account double and triple simultaneous steps.

As an illustration of an application, we implemented our tracking algorithm to measure the unfolding of native chromatin, shown in Fig 5d (experimental details in [53]). Native chromatin is an example of a highly heterogeneous sample, where many molecules need to be measured to extract common features. We used 3DPT to measure the step size of the unwrapping of DNA from the histone core, using the method of the step size described by Kaczmarczyk [56] and summarized in Materials & Methods. As expected, 3DPT accurately revealed the characteristic 77 bp step size, corresponding to about 25 nm, shown in figure 5e. Although the molecules were heterogeneous in composition and unfolded accordingly, the width of the individual steps occurring at forces above 5 pN could easily be resolved.

## Discussion and Conclusion

Single-molecule biophysical techniques frequently employ bead tracking for the mechanical characterization of biomolecules. Here, we introduced 3DPT, an efficient and non-iterative bead tracking algorithm for holographic imaging. It makes use of the circular symmetry of a holographic image of a bead and, similar to lock-in techniques, it selects a single spatial frequency for image analysis. Next to directly producing the *xy*-position, the height information is captured in a single parameter: the phase of the wave front of the diffraction pattern, which can be converted to a *z*-coordinate using calibration prior to the experiment. 3DPT was lightweight, robust against common aberrations, and yielded nanometer accuracy in three dimensions. We have implemented this algorithm in NI LabVIEW 2014 on a 10 core 2.8 GHz CPU and could track up to 10.000 diffraction patterns per second captured within a ROI of 100×100 pixels. 3DPT is robust against many common image artifacts, and can yield nanometer accuracy despite sub-optimal imaging conditions

Due to drift and mechanical vibrations in our setup, we did not test whether 3DPT could resolve nanometer steps, as was demonstrated before [51]. Nevertheless, 25 nm steps, resulting from unwrapping single nucleosome were easily resolved. From PSD analysis, as well as unfiltered time traces, it is clear that 1 nm is close to the limit of the accuracy of our current experiments at 30 Hz bandwidth. 1 Hz averaging should be able to resolve nanometer changes in bead position. In the lateral direction, 3DTP performs an order of magnitude better. For many applications other than single-molecule force spectroscopy, the lateral resolution is as important as accuracy in the *z*-direction.

Computing the radial intensity profile is the most time-consuming step in traditional bead tracking algorithms and transferring this step to GPU can increase the processing speed [50]. We nevertheless do not expect a significant gain in speed by implementing 3DPT in GPU, as the current computation times are smaller than the time it takes for transferring the images from CPU to GPU. Avoiding computation of the radial intensity profile not only increases the speed, but it also makes the tracking more robust: small errors in *xy*-position, as well as imaging aberrations, introduce large changes in this profile as compared to a reference image. In 3DPT, these artifacts predominantly reduce the amplitude of the cross-correlation, but have little effect on the position or the phase.

Because the computer-generated reference images that are used in 3DPT have a known center and lack noise and other artifacts, the algorithm is very robust. We therefore expect that 3DPT may be used for other applications than the tracking of spherical colloidal particles. This, however, should be tested for each particular application. In addition to bead tracking, we also used a cross-correlation with reference images for auto-focusing and for initial recognition of beads, relieving the operator from manually selecting beads. For multiplexed high-throughput applications, this can be a key advantage as the microscopy can be fully automated.

We could not obtain good fits of our super-paramagnetic beads to LMST, resulting in large tracking errors, typically hundreds of nanometers, which we tentatively attributed to the heterogeneous composition of the beads. This suggests that LMST fitting is a more viable option for, for example, TPM, AFS, and OT, that do not require magnetic beads. Though fitting imprecisions did not significantly affect the resulting position accuracy in the *xy*-direction, it was detrimental for tracking in the *z*-direction. Fits only converged in a limited range, especially in the case of the 2.8 *μm* beads. For MT, LMST fitting therefore may not only impede real-time processing, but it may also be inadequate for applications that require more than several μm range.

In previous work, the Grier group used a laser to create holographic images [41]. Most biophysical single-molecule studies (including ours), however, used a collimated LED to illuminate the sample. Due to the limited spatial coherence of a LED, the images do not feature speckle patterns, that were subtracted in studies using LMST fitting [61]. The limited coherence also reduces the range of the diffraction pattern, and laser illumination in combination with image background subtraction may further improve the accuracy of 3DPT by generating more contrast in the holographic image.

Multiplexing becomes increasingly important in single-molecule biophysics as more complex and more heterogeneous samples are investigated. A good example is our previous study, in which we performed force spectroscopy on natively assemble chromatin fibers [53]. Multiplexing allowed us to pick out hundreds of chromatin fibers that each unfolded differently, reflecting variations in composition and folding. Another example is a study on RNA polymerase pausing, in which multiplexing served to observe sufficient rare events to analyze the statistics [62]. Because 3DPT can easily replace traditional radial profile comparisons with LUTs, it may be adopted by many experimentalists.

Overall, its easy implementation, robustness, and nanometer accuracy over a wide *z*-range makes 3DPT an ideal method for multiplexed particle tracking applications. It may improve all single-molecule techniques that rely on bead tracking such as MT, TPM, AFS, and OT. By enhancing their throughput, it will help to realize one of the truly unique promises of single-molecule biophysics: detecting rare events in a large population of molecules with unprecedented resolution.

## Author Contributions

T. B. B. and J. v. N. conceived the study. T. B. B., N. H. and J. v. N. designed and performed the research. T. B. B. and J. v. N. analyzed data, performed the simulations, and developed the mathematical framework. N. H. measured and analyzed the chromatin data.

## Acknowledgments

The authors would like to thank Artur Kaczmarczyk for valuable discussions during the development of the algorithm. This work is part of the research program VICI with project number 680-47-616, which is (partly) financed by the Netherlands Organization for Scientific Research.

---

**Figure S1.**
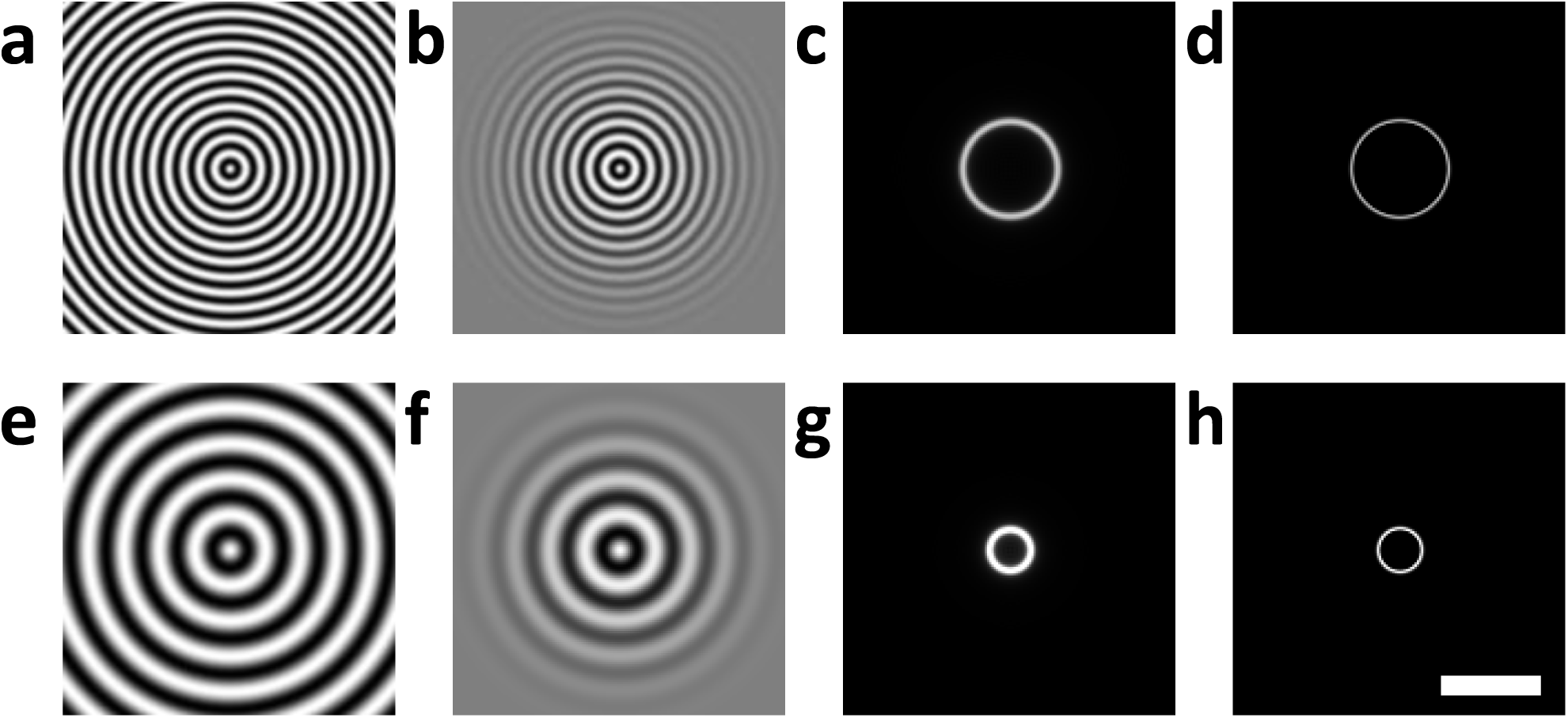

